# Inhibition of H3K27M-enhanced ATM signaling increases radiation efficacy in diffuse midline glioma

**DOI:** 10.1101/2024.11.01.621526

**Authors:** Erik Peterson, Leslie A. Parsels, Joshua D. Parsels, Xinyi Zhao, Maria G. Castro, Theodore S. Lawrence, Carl Koschmann, Qiang Zhang, Daniel R. Wahl, Meredith A. Morgan

## Abstract

H3K27-altered diffuse midline glioma (DMG) is an aggressive and treatment-resistant form of pediatric high-grade glioma (pHGG). The disease is defined by point mutations in histone H3 that convert lysine 27 to methionine (termed H3K27M), resulting in genome-wide epigenetic changes that drive tumorigenesis. While radiation therapy is the standard of care, subsequent recurrence, often within the high dose radiation field, is universal. We found that the apical DNA damage response (DDR) kinase Ataxia Telangiectasia-Mutated (ATM) was uniquely upregulated in H3K27M-expressing patient tumor samples compared to pHGG expressing only wild-type histone H3. Using a panel of H3K27 isogenic DMG cell lines, we further found that H3K27M was associated with reduced H3K27me^3^ within the *ATM* promoter, increased ATM mRNA levels, and elevated DDR signaling, even in the absence of exogenous DNA damage. Consistent with these results, AZD1390, a clinical-grade, CNS-penetrant ATM inhibitor, sensitized H3K27M neurospheres to the long-term effects of radiation on survival, in part due to attenuated repair of radiation-induced DNA damage. Finally, AZD1390 sensitized orthotopic H3K27M mutant tumors to radiotherapy and significantly extended median survival relative to vehicle, AZD1390 or radiation alone (50 days vs 31, 36 or 39 days, respectively) with minimal adverse effects. Taken together, these data provide a direct mechanistic link between the H3K27M mutation and ATM expression and support the clinical investigation of AZD1390 with radiotherapy in H3K27M-altered DMG.

## INTRODUCTION

Diffuse midline glioma (DMG) is an aggressive and uniformly fatal pediatric high-grade glioma (pHGG) with a median survival of approximately one year (1–3). Fractionated radiation therapy is the standard of care for these tumors. Moreover, they derive little benefit from chemotherapy and typically cannot be surgically removed due to their midline location (1). Unfortunately, DMG inevitably recurs within the high dose radiation field (2). As toxicity to surrounding normal tissues prevents further escalation of the radiation dose (3), there is a critical need to develop selective radiosensitizing strategies for DMG that enhance the therapeutic efficacy of radiation without increasing neurotoxicity.

DMG is defined not only by anatomical location, but also by monoallelic point mutations that change the 27^th^ residue of tail domain of histone H3 from lysine to methionine (termed H3K27M)(4,5). Although this mutation is present in only a small percentage of the total histone H3 pool, H3K27M-mediated suppression of polycomb repressive complex 2 (PRC2) activity leads to distinct, widespread epigenetic changes, including global chromosomal hypomethylation of H3K27, decompaction of chromatin, and the activation of aberrant transcriptional profiles that promote tumorigenesis and have been hypothesized to affect therapeutic response (4–7).

While radiation (RT) induces numerous types of DNA damage, unrepaired DNA double-strand breaks (DSBs) are the lethal lesions responsible for tumor cell killing (8). These lesions activate a network of intracellular signaling pathways that regulate cell cycle progression, DNA replication and repair, and/or apoptosis and are collectively termed the DNA damage response (DDR). Small molecule inhibitors of the DDR increase the efficacy of radiation therapy across several cancer types, including pHGG (9–13). While the combination of DDR inhibitors with RT is effective in preclinical models of DMG (14), whether the H3K27M mutation specifically affects the DNA damage response and/or offers a tumor cell-selective target for DNA damage response inhibitors in combination with RT were questions that remained unknown.

In this study, we found that both H3K27M-expressing DMG patient tumors and patient-derived cell lines preferentially upregulate the expression of ATM, one of the three apical phosphytidal-inositol-3-kinase-like DDR proteins that play a vital role in the repair of RT-induced DSBs. Furthermore, we found reduced H3K27 trimethylation (H3K27me^3^) in the *ATM* promoter of H3K27M mutant DMG cells relative to matched isogenic control cells that was associated with increased ATM expression and signaling. This altered regulation of ATM suggested that ATM inhibition would increase the sensitivity of H3K27M DMG cells to radiation, an observation also seen by other groups (9). To test this hypothesis, we assessed the ability of AZD1390, a CNS-penetrant ATM inhibitor, to radiosensitize isogenic H3K27M mutant and wildtype DMG cells. Subsequently, we tested the ability of AZD1390 to radiosensitize *in vivo* in orthotopic H3K27M mutant DMG tumors, including molecular correlate studies to assess ATM inhibition and DNA damage in both tumor and normal brain. Collectively, the results of this study establish a unique mechanistic connection between H3K27M, ATM expression, and DDR signaling, as well as support development of AZD1390 with RT as a therapeutic strategy for H3K27M DMG.

## METHODS

### Cell Lines and Reagents

Both the human PDX BT245 and DIPG.XIII H3K27M-knock out isogenic models and the murine PPW/PPK H3K27M isogenic model have been previously reported (15,16). Human cells were maintained as neurosphere (suspension) cultures in 1:1 Neurobasal-A Medium:DMEM/F-12, GlutaMAX Medium supplemented with 10 mM HEPES [4-(2-hydroxyethyl)-1-piperazine ethanesulfonic acid], 1 mM sodium pyruvate, 0.1 mM MEM non-essential amino acids, 2 mM GlutaMAX, and antibiotic-antimycotic (all Gibco/Life Technologies); B27-A supplement (without Vitamin A) (50X, Invitrogen); heparin (2 µg/ml; Stemcell Technologies, Inc.); 20 ng/mL human EGF and FGFb and 10 ng/mL platelet derived growth factors AA and BB (PeproTech) as previously described (17). The murine PPW and PPK isogenic cell lines were established from tumors derived by *in utero* electroporation of plasmids expressing dominant-negative TP53, mutant PDGFRA (D842V) and either wild-type histone H3 (H3WT) or mutant histone H3K27M, respectively, and were cultured in Neurobasal-A Medium supplemented with B27-A with Vitamin A (50X, Invitrogen), N2-supplement (100x, Invitrogen), 1 mM sodium pyruvate, 0.1 mM MEM non-essential amino acids, 2 mM L-glutamine, antibiotic-antimycotic and 20 ng/mL human EGF as previously reported (15). DIPGXIII.K27M cells expressing both GFP and firefly luciferase (DIPG.XIII.GFP/LUC) were established by lentiviral infection with pLenti-GF1-CMV-VSVG (University of Michigan Vector Core; product discontinued) into parental DIPG.XIII cells. AZD1390 (AstraZeneca) was obtained directly from the supplier. For *in vitro* studies, AZD1390 was dissolved in dimethyl sulfoxide and stored in aliquots at −20 C. For *in vivo* studies, a suspension of 5 mg/mL AZD1390 was prepared in 0.5% w/v hydroxypropyl methylcellulose (HPMC) and 0.1% w/v Tween 80.

### Tumor Expression Data

mRNA-seq Z-score expression data originally reported by Mackay et al. were obtained through PedCBioPortal (Children’s Hospital of Philadelphia Research Initiative)(18). Z-scores for each gene of interest were extracted and sorted based on histone H3 mutational status (either wild-type or H3K27M-mutant histone H3 (*H3F3A* or *HIST1H3B*)). Only patient samples with confirmed histone H3 mutational status were used.

### Quantitative RT-PCR

Total RNA was isolated using the RNeasy Kit per manufacturer’s protocol (Qiagen). cDNA was prepared using the High-Capacity Reverse Transcriptase Kit (Applied Biosystems) with oligo dT primers. Quantitative PCR was performed using PowerTrack™ SYBR Green Master Mix (ThermoFisher) on a QuantStudio 6 Flex Real Time PCR System (Applied Biosystems). Expression data were analyzed using the ΔΔCt method against β-actin (housekeeping control) and were normalized to matched H3K27M-wt or KO isogenic samples. Primer sequences are listed in Supplementary Data (**Table S1**).

### ChIP-PCR

Chromatin immunoprecipitation (ChIP) was performed using the SimpleChIP® Enzymatic Chromatin Magnetic Bead IP Kit per manufacturer’s protocol (Cell Signaling #9003). At least 4×10^6^cells were plated in quadruplicate in 10 cm dishes and grown for ∼2 days until they began to form small spheres. Nuclear proteins were crosslinked to DNA with formaldehyde, replicate samples were pooled, and nuclei were isolated and digested with micrococcal nuclease for 45 min at 37°C. Digested nuclei were pelleted and resuspended in ChIP buffer and sonicated with five 20 second pulses interspersed with 30 seconds on wet ice. Lysates were then clarified, and chromatin concentration was quantified and normalized to 5-10 μg of DNA per IP reaction. Antibodies were added to the DNA, which was incubated overnight at 4°C on a rotator. Samples were incubated with magnetic beads which were then placed in a magnetic tube rack to allow for removal of the supernatant. Beads were then washed several times before chromatin was eluted, de-crosslinked, and purified. Primer sequences and antibodies used are listed in Supplementary Data (**Tables S1** and **S2**, respectively). Enrichment is expressed as the Percent of Input (% Input) and data are normalized to matched H3K27M-wt or KO isogenic cells.

### Irradiation

Both *in vitro* and *in vivo* radiation studies were performed using a Philips RT250 (Kimtron Medical) at a dose rate of approximately 2 Gy/min at the University of Michigan Rogel Comprehensive Cancer Center Experimental Irradiation Shared Resource (Ann Arbor, MI). Dosimetry was performed using an ionization chamber connected to an electrometer system that is directly traceable to a National Institute of Standards and Technology calibration. For tumor irradiation, animals were anesthetized with isoflurane and positioned in pairs such that the entire cranium of each mouse was at the center of a 2.4 cm aperture in the secondary collimator, with the rest of the mouse shielded from radiation.

### Western Blotting

Whole cell lysates were prepared in ice-cold RIPA buffer (1M NaCl, 1% NP-40, 0.5% sodium deoxycholate, 0.1% SDS, 50 mM Tris, pH 7.4) supplemented with both PhosSTOP^™^ phosphatase inhibitor (Roche) and cOmplete^™^ Protease Inhibitor Cocktail (Roche). After heat-denaturing at 95°C for 10 minutes, samples were separated by SDS-PAGE and transferred to Immobilon^®^-P PVDF Membrane (0.45mm pore size; Millipore) per standard protocols. Membranes were blocked with 5% milk diluted in tris-buffered saline/0.075% Tween20 (TBST) for 30 min at room temperature, rinsed with TBST and then incubated with primary antibodies diluted in 5% bovine serum albumin (Millipore Sigma A3912)/TBST overnight at 4°C. Membranes were incubated with secondary antibodies for 1 h at room temperature, washed, and developed in either SuperSignal™ West Dura Extended Duration Substrate (Thermo Scientific) or BrightStar™ Duration HRP Chemiluminescent Substrate (Alkali Scientific). For tissue analyses, bulk tumor tissue was homogenized in RIPA buffer containing protease and phosphatase inhibitors using a tissue homogenizer (FisherBrand). Protein concentration was measured using the Pierce™ BCA Protein Assay Kit (ThermoFisher). Primary antibodies used are listed in Supplementary Data (**Table S2**).

### Neutral Comet Assay

H3K27M-isogenic cell lines were plated in 6 well plates and allowed to grow for ∼2 day until they became small spheres. One hour before radiation, cells were treated with vehicle control or 300 nM AZD1390 and incubated at 37°C. Treated cells were irradiated with a single 8 Gy dose of radiation as previously described and immediately returned to a 37°C incubator. At the appropriate time points, cells were collected and centrifuged at 4°C before being placed on ice. Cells were then trypsinized and passed through a 40μm strainers to ensure single cell suspension in cold PBS. Cells were then added to 1% low melting point agarose at low density (10,000 cells/mL) and 50μL of suspension was added to ice-cold CometSlides (R&D Systems #3042135) on wet ice in a lidded cooler. Slides were kept covered to prevent any potential DNA damage from visible light. Slides were lysed overnight in Comet Assay Lysis Solution (R&D Systems). The next day, DNA was electrophoresed in a CometAssay ESII tank (Bio-techne), and DNA was precipitated using DNA precipitation buffer at room temperature. Slides were then washed and dried briefly at 37°C before being stained with 2.5μg/mL propidium iodide. Slides were then rinsed and dried at 37°C. Before imaging, 50μL of PBS was added to each sample before coverslips were placed. Images were captured using a Nikon Eclipse Ti2 microscope with attached DS-Ri2 camera using a red fluorescence filter. Comet tails and olive tail moments (OTM) were measured for a minimum of 50 cells per sample using Comet Assay IV software (Instem).

#### CellTiter-Glo® 3D Cell Viability Assay

PPW / PPK, BT245 and DIPG.XIII isogenic cells were plated into 6 well dishes (2×10^5^ cells/well) and allowed to grow into small neurospheres prior to treatment (∼2 days). AZD1390 was given for 25 h, beginning 1 h prior to RT. After treatment, spheres were dissociated with Accutase (StemCell Technologies), passed through a 40μm strainers and plated in quadruplicate as single cell suspensions in 96 well plates (Corning Costar #3610). Plated cells were imaged at t_0_ to record the exact number of cells per well. Five days later, plates were analyzed with the CellTiter-Glo® 3D Cell Viability Assay Kit (Promega), which uses ATP levels as a surrogate for viability, per manufacturer’s protocol. Luminescence values for each well were normalized to the number of cells originally plated in that well. Radiation survival was normalized for drug toxicity and the radiation enhancement ratio (RER) for a given dose of radiation was calculated as the ratio of the survival under control conditions divided by the survival after drug exposure. A value significantly greater than 1 indicates radiosensitization. Cytotoxicity in the absence of radiation treatment was calculated by normalizing the normalized luminescence of drug treated-cells to non-drug treated cells.

#### Long-term Neurosphere Growth Assay

Cells treated as described in *CellTiter-Glo® 3D Cell Viability Assay* were plated in quadruplicate as single cell suspensions in 96 well plates (Corning Costar #3610) which were transferred to a BioTek BioSpa 8 Automated Incubator (Agilent Technologies) and imaged with the 4x objective every 12 h for 10-14 days using a BioTek Cytation 5 Cell Imaging Multimode Reader (Agilent). Images were processed for data acquisition using BioTek GenV software (Agilent). Each well was imaged at t_0_ to record the exact number of cells seeded per well. Neurospheres with an area larger than 2,600 µm^2^ were scored as surviving cells. The endpoint of each experiment was determined by when the number of spheres began to decline due to sphere merging. Plating efficiency (PE), determined by the number of surviving spheres normalized to the number of cells at t_0_, was monitored over time for each condition, and the PE at the time a sample reached its growth plateau was used to calculate the surviving fraction (SF; see Supplementary Data Fig. S1). Radiation survival was normalized for drug toxicity and the RER was calculated as the ratio of the radiation survival under control conditions divided by the radiation survival after drug exposure.

### Immunofluorescence

For immunofluorescence experiments, DIPG.XIII isogenic cells were grown as described in *Survival Assays*. Following treatment and accutase dissociation of neurospheres, cells were fixed with an ice-cold solution of 3.7% paraformaldehyde, 2% sucrose, 0.5% Triton X-100 in PBS. Samples were stained with a mouse monoclonal γH2AX antibody (JBW301, Millipore Sigma) and DAPI (4’6-diamidino-2-phenylindole) as previously described (19). Samples were then washed and resuspended in 0.2% Triton X-100/0.5% BSA diluted in ddH_2_0 and spotted on to positively charged glass slides (20). After drying, coverslips were mounted with ProLong™ Gold Antifade (ThermoFisher) and samples were visualized with an Olympus IX73 microscope (Olympus America) using the 60x oil objective. Fields to score were chosen at random based on DAPI staining. Cells with 10 or more γH2AX foci were scored as positive. At least 100 cells from each of 2-4 independent experiments were manually scored for each condition.

### Stereotactic Orthotopic Implantation

Animals were handled according to a protocol approved by the University of Michigan Committee for Use and Care of animals. Rag1-KO C57BL/6 mice were anesthetized with ketamine and administered carprofen analgesic before removing the scalp fur and sterilizing the incision site. Mice were then placed into a stereotactic injection rig to prevent movement during the procedure. An incision was made along the midline of the scalp and a small hole in the skull was made using an electric burr hole drill at coordinates 3 mm lateral, 1 mm rostral from the bregma. Approximately 1.5 – 2.0 ×0^5^ cells in a volume of 3 μL (∼5.0×10^4^ cells/μL) were implanted in the cortex at depth of approximately 2.5 mm using 10 μL Hamilton syringes. After injections, wounds were sutured and treated with triple antibiotic ointment. Mice were given atipamezole to reverse the anesthetic and placed in heated cages during recovery from anesthesia. Mice were provided diet gel supplement and monitored for 10 days, including a second dose of carprofen the day following the surgery.

### Bioluminescent Imaging and Mouse Treatments

Mice were administered sterile filtered 30 mg/mL luciferin solution (Syd Laboratories) via IP injection. Ten minutes post-injection, bioluminescence (BL) was measured using an IVIS Spectrum imaging station (University of Michigan Imaging Core). Bioluminescence flux values were used to randomize mice into four treatment groups (8 mice/group) so that the average bioluminescent flux was similar between groups. Imaging was repeated 1-2 times weekly during efficacy studies. Treatment groups included: vehicle control (0.5% w/v HPMC, 0.1% w/v Tween 80), AZD1390 alone, RT alone, and AZD1390 + RT. AZD1390 (20mg/kg) was administered via oral gavage following the schedule of three days on, two days off, and three days on. For radiation treatment, mice were anesthetized using 2.5% isoflurane in an induction chamber and via nose cones within the irradiation chamber. Mice were positioned such that the head was placed under a flat collimator with a lead shield covering the entirety of the body except for the whole brain. Mice were administered a total of six 2 Gy doses of radiation, 1 hr after AZD1390 treatment, on a schedule of three days on, two days off, three days on. Mice were euthanized upon development of neurological symptoms.

### Immunohistochemistry

Whole brains were collected from mice and tumors were identified using GFP-fluorescence guidance. The brain was bisected along the mid-tumor line and one half of the brain was saved for IHC. Formalin-fixed, paraffin-embedded tissue samples were cut into slides using a Leica HistoCore BIOCUT microtome. Slides were baked overnight in a 56°C incubator, then deparaffinized and subjected to antigen retrieval. Slides were blocked with goat serum and stained with primary antibody overnight. Antibodies used are listed in Supplementary Data (**Table S2**). Tissues were then stained with 3,3’-diaminobenzidine (DAB). Stained slides were incubated in hematoxylin, 1% clarifying reagent, and bluing agent before being washed in ethanol and xylene and sealed with neutral gum. Slides images were captured with a Nikon Eclipse Ti2 microscope with attached DS-Ri2 camera. Images were processed using NIS-Elements (Nikon) software. DAB staining quantification was performed on at least 500 cells per tissue slice using QuPath 0.4.3 software.

### Statistics

All data are presented as the mean ± SEM unless otherwise stated. When assessing statistical significance between treatment groups, continuous variables were analyzed using the unpaired Student t-test, and Student t-test with Welch’s correction, for normally and non-normally distributed data, respectively. Mouse survival data were analyzed using the Kaplan-Meier method and log-rank test. P-values <0.05 were considered significant and are denoted in the figures as follows: *p<0.05, **p<0.01, ***p<0.001. Statistical analyses were performed in GraphPad Prism 10.3.0 (GraphPad Inc.).

## RESULTS

### H3K27M mutation increases *ATM* expression and ATM signaling

The H3K27M mutation fundamentally alters the DMG cellular epigenome, driving aberrant gene expression (4,5,21,22). We hypothesized that there are H3K27M-driven epigenetic changes in the DNA damage response that we may be able to target pharmacologically to selectively sensitize H3K27M mutant DMG tumors to radiotherapy. We first assessed differences in the expression of DDR genes in pediatric high-grade glioma (pHGG) samples from patient tumors expressing either wild-type histone H3 (H3WT) or the H3K27M mutant protein. Using publicly available mRNA-seq Z-score data (18), we determined that pHGG tumors expressing H3K27M had significantly higher expression of *ATM* than those expressing only the histone H3 wild-type protein (**Fig. 1A**). Further, there were no H3K27M-associated differences in the expression of other DDR genes with potential clinical relevance for DMG, such as *ATR*, *PRKDC*, *PARP1*, *WEE1* or *CHEK1* (11,19,23–25). These results suggest that among the DDR genes, *ATM* is uniquely upregulated in H2K27M-expressing pHGG.

**Figure 1:**
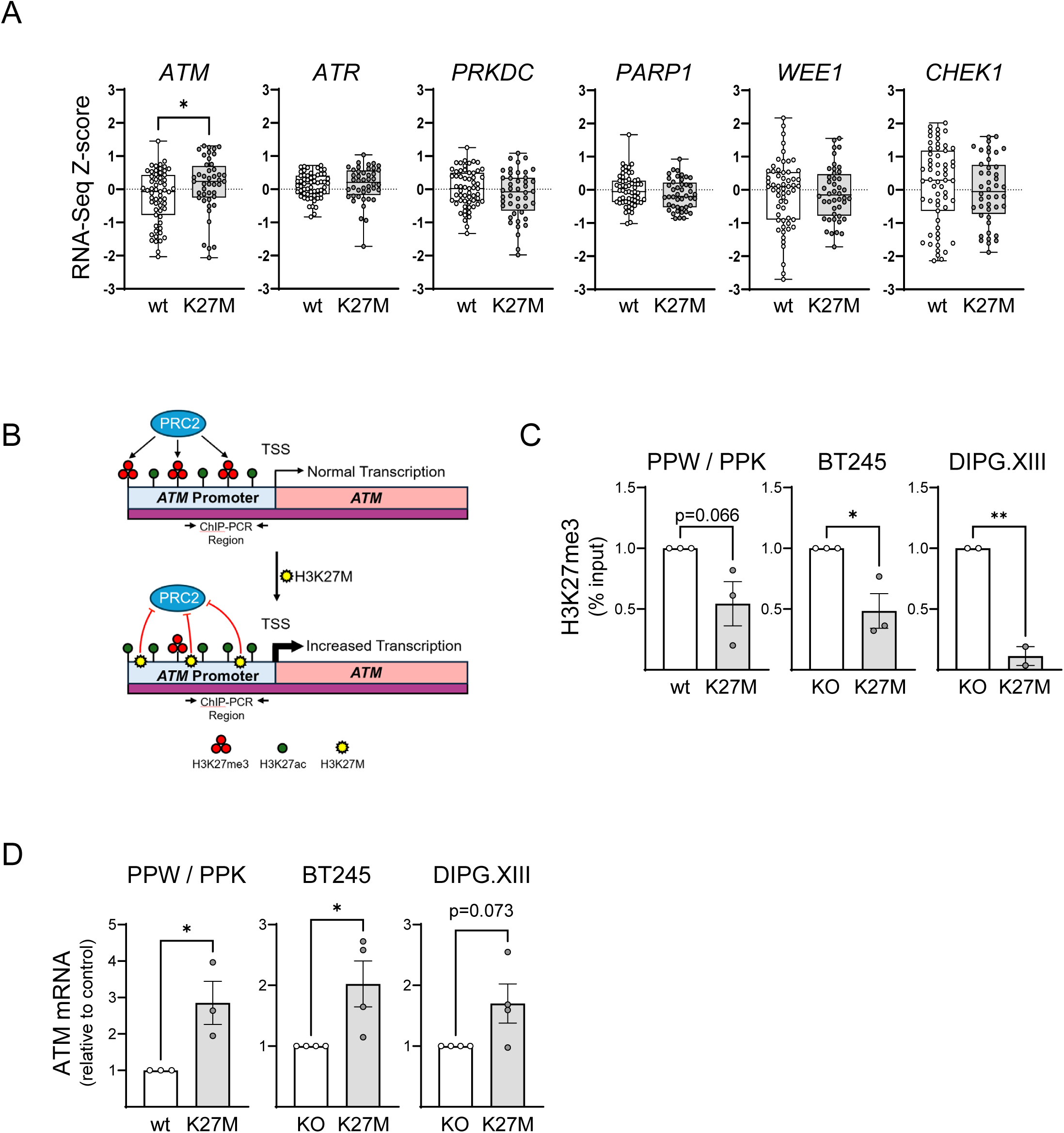
Expression of mutant histone H3K27M is associated elevated expression of *ATM*. (**A**) RNASeq Z-score analyses of publicly available data (PedcBioPortal) for relative expression of DNA damage response genes in pediatric high-grade gliomas expressing either wild-type histone H3 (H3WT) or K27M-mutant histone H3. (**B**) Model of H3K27M-associated epigenetic changes in the ATM promotor region that have the potential to upregulate transcription of *ATM* (PRC2: polycomb repressive complex 2; TSS: transcriptional start site). (**C**) ChIP analysis of the repressive epigenetic mark H3K27me^3^ at the ATM promotor. (**D**) qRT-PCR analysis (ddCT) of relative ATM expression levels in isogenic models of histone H3K27M loss. For primer sequences and antibodies used for ChIP analysis, see Supplementary Data Tables S1 and S2, respectively.

To determine whether this increase in ATM expression resulted from H3K27M-driven epigenetic changes affecting transcriptional regulation of *ATM,* or was part of a broader response to the genetic instability associated with H3K27M mutation in DMG (26), we used previously described isogenic histone H3WT and H3K27M cell lines (15,16) to determine whether H3K27M directly affects *ATM* expression. One of the primary epigenetic changes associated with H3K27M is a global reduction in levels of the transcriptional repressor H3K27me^3^ (**Fig. 1B**) (16). We found that H3K27M-expressing cells had reduced levels of H3K27me^3^ specifically within the *ATM* promotor region (**Fig. 1C**) that correlated with higher *ATM* transcript levels (**Fig. 1D**).

To determine whether these transcriptional and epigenetic changes had downstream biologic effects, we interrogated ATM signaling and protein levels. While there was a modest trend towards increases ATM protein in the H3K27M mutant cells, constitutive DDR signaling, including the ATM autophosphorylation site S1981 and the downstream effector proteins pKAP1 (S824)(27), pCHK2 (T68), and pCHK1 (S345), was significantly higher in the H3K27M-expressing cells (**Fig. 2A, B**). This increase in DDR signaling was associated with both a higher level of innate DSBs, as determined by neutral comet assay, and a trend towards higher induction of DSBs immediately following high dose radiation (**Fig. 2C, D**). Collectively, these results support the hypothesis that H3K27M mutation reduces H3K27me^3^ within the ATM promoter, leading to enhanced ATM expression and signaling that are associated with elevated levels of DNA damage in H3K27M mutant DMG.

**Figure 2:**
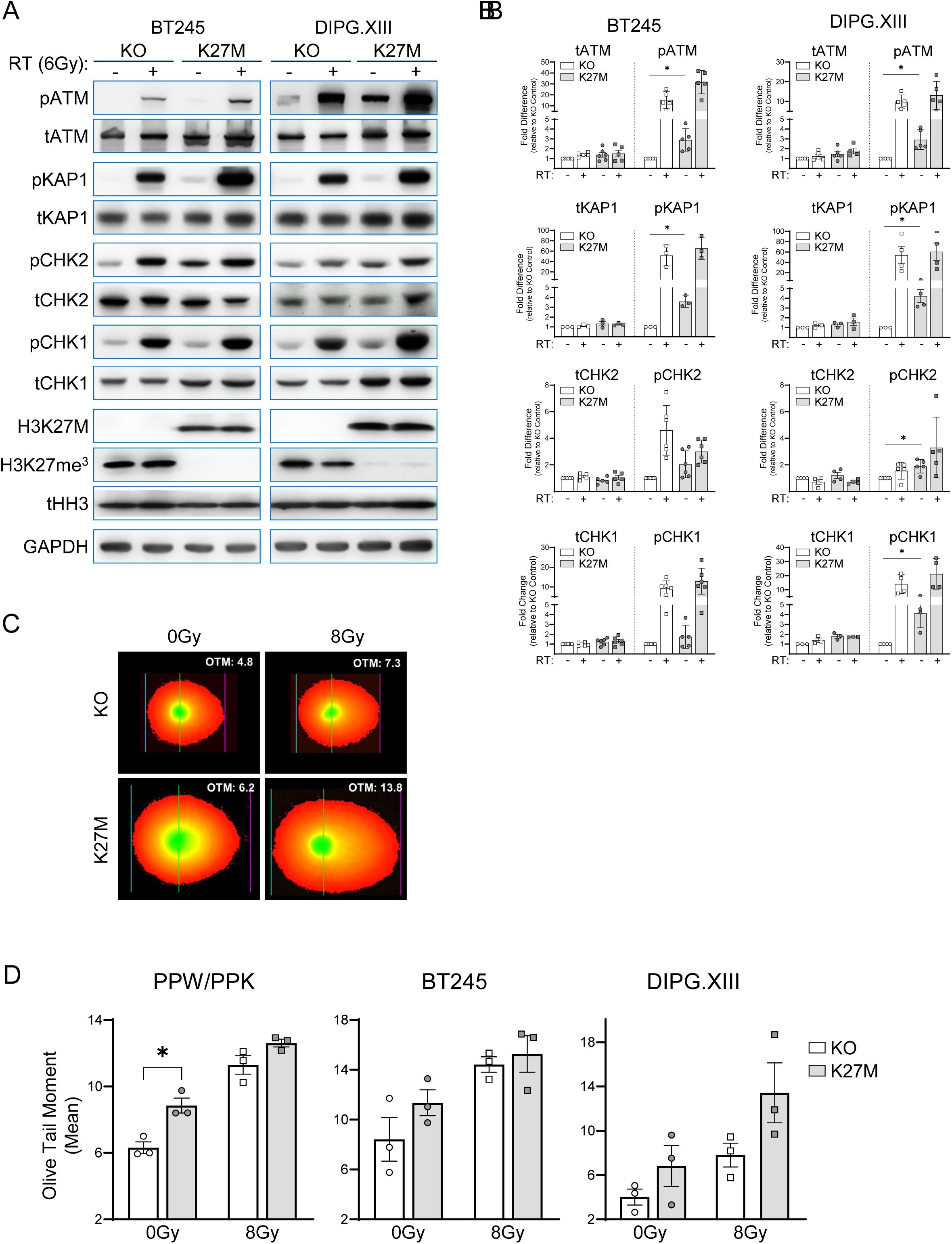
Increased ATM signaling and DNA damage in H3K27M mutant cells. (**A**) Immunoblots of isogenic cells under baseline conditions or at 2 h post-RT (6 Gy). Images are representative of results from 3 independent experiments. (**B**) Quantitation of total and phospho-proteins shown in panel A, normalized to respective H3K27 wildtype condition. Data are the mean ± SEM of 3-6 independent experiments. (**C**) DSB levels in H3K27 wildtype versus H3K27M mutant cells measured by neutral comet assay under control conditions or 2 h post-RT (8 Gy). Data are the mean ± SEM of 3 independent experiments. (**D**) Representative images of DIPGXIII comets.

### ATM inhibition sensitizes H3K27 mutant DMG cells and neurospheres to radiation

Targeting the DDR to increase radiosensitivity has been used in a variety of cancer types, including adult glioma and DMG (28). To determine whether AZD1390 radiosensitized glioma cells in an H3K27M-dependent manner, we first assessed the effects of AZD1390-mediated ATM inhibition on the radiosensitivity of isogenic histone H3WT and H3K27M cell lines using the CellTiter-Glo 3D viability assay (**Fig. 3A-C**). In the absence of ATM inhibitor, we found similar baseline radiation sensitivity between H3K27 wild-type or knockout and mutant PPK and BT245 cells, but H3K27M-associated radiation resistance (6Gy) in mutant DIPGXIII cells (**Suppl. Table 3**). Furthermore, while AZD1390 significantly radiosensitized in all six DMG cell lines, only in the PPW/PPK model did H3K27M mutation confer greater sensitivity to AZD1390-mediated radiosensitization (though this difference did not reach statistical significance). While these viability assays indicated radiosensitization by AZD1390, we sought to further evaluate radiosensitization by assessment of RT-induced loss of clonogenicity using long-term sphere survival assays. Given the clinical significance of the H3K27M mutation in DMG, as well as the amenability of H3K27M mutant cells to neurosphere survival assays, H3K27M mutant neurospheres were treated with AZD1390 and increasing doses of radiation followed by dissociation, replating as single cells, and live-cell imaging to monitor sphere formation and survival over time (**Suppl. Fig. S1**). Surviving fractions were calculated from the ‘plateau’ plating efficiencies, when the number of mature spheres (spheres >2600 μm^2^) first plateaued, for each condition. We found that AZD1390 significantly reduced neurosphere survival following radiation in both PPK and BT245 cells (**Fig. 3D, F**), with 2 Gy radiation enhancement ratios of 1.63±0.10 and 1.82±0.16, respectively (**Fig. 3E, G; Suppl. Table 3**). While the results generally show concordance between the viability and long-term survival assays, there was a significant difference in DIPG.XIII radiosensitivity determined by CellTiter-Glo (6Gy SF=1.01±0.05) compared to that determined by neurosphere survival (**Fig. 3H**; 6Gy SF=0.57±0.01; *p<0.05); a difference that likely reflects the variable growth rate of these cells over time (**Suppl. Fig. S1).** Despite these differences, AZD1390 radiosensitized DIPGXIII cells in both assays although to a lesser extent than in other cell lines (**Fig. 3I**). Taken together, these results confirm that AZD1390 is an effective radiosensitizing agent in preclinical models of DMG.

**Figure 3:**
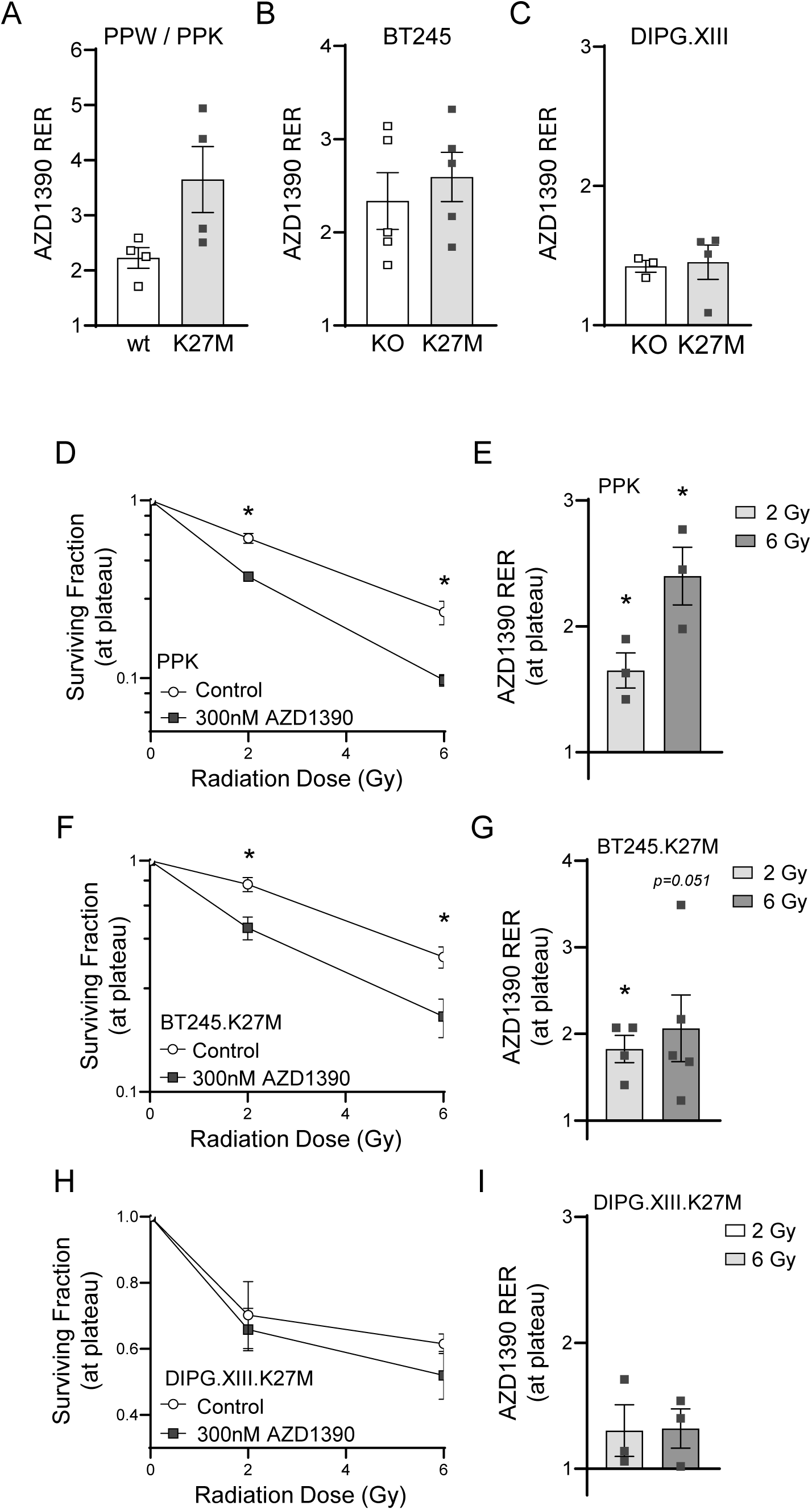
Inhibition of ATM radiosensitizes H3K27M mutant cells and neurospheres. (**A-C**) Cell proliferation assays were used to assess radiosensitization of DMG cells treated with minimally cytotoxic concentrations of AZD1390 (300 nM) and 2 Gy (PPW/PPK, BT245) or 6 Gy RT (DIPGXIII). See also, Supplementary Data, Table S3. (**D-I**) Neurosphere survival and radiosensitization by AZD1390 were assessed by long-term neurosphere growth assays. Survival data were determined based on the plating efficiency calculated at the growth plateau for each condition (see Supplementary Data, Fig. S1). Data are the mean ± SEM from 3-5 independent experiments (*p<0.05 vs. survival under control conditions; **D, F, H**). Radiation enhancement ratios were calculated from the corresponding survival data (*p<0.05 vs. control; **E, G, I**). See also, Supplementary Data, Table S3.

### ATM inhibition reduces DDR signaling and increases RT-induced DNA damage in H3K27M DMG

ATM autophosphorylation is an initiating event in the response to and repair of radiation-induced DSBs (29). Furthermore, the persistence of radiation-induced DNA damage is a hallmark of DDR inhibitor-mediated radiosensitization. To assess the ability of AZD1390 to inhibit ATM activity and its downstream signaling in DMG cells, we evaluated the phosphorylated forms of ATM and downstream DDR effector proteins under both control conditions and at 2 h post-RT (6 Gy). As expected, we found that AZD1390 inhibited baseline autophosphorylation of ATM at S1981 (pATM-S1981), as well as induction pATM-S1981 in response to RT, in both H3WT and H3K27M cells (**Fig. 4A, B**). AZD1390 treatment also inhibited radiation-induced phosphorylation of the ATM effector protein KAP1 (pKAP1-S824)(27), though that effect was mildly attenuated in the H3K27M cells, possibly due to the elevated ATM activity in those cells. In contrast, while AZD1390 significantly reduced constitutive CHK2 phosphorylation (pCHK2-T68) in both H3WT and H3K27M cells, it had little effect on RT-mediated phosphorylation of T68-CHK2. This difference likely reflects redundant RT-mediated DDR kinase activity upstream of CHK2. Regardless, AZD1390 effectively inhibited ATM autophosphorylation and downstream signaling in both H3K27 wild-type and mutant cells.

**Figure 4:**
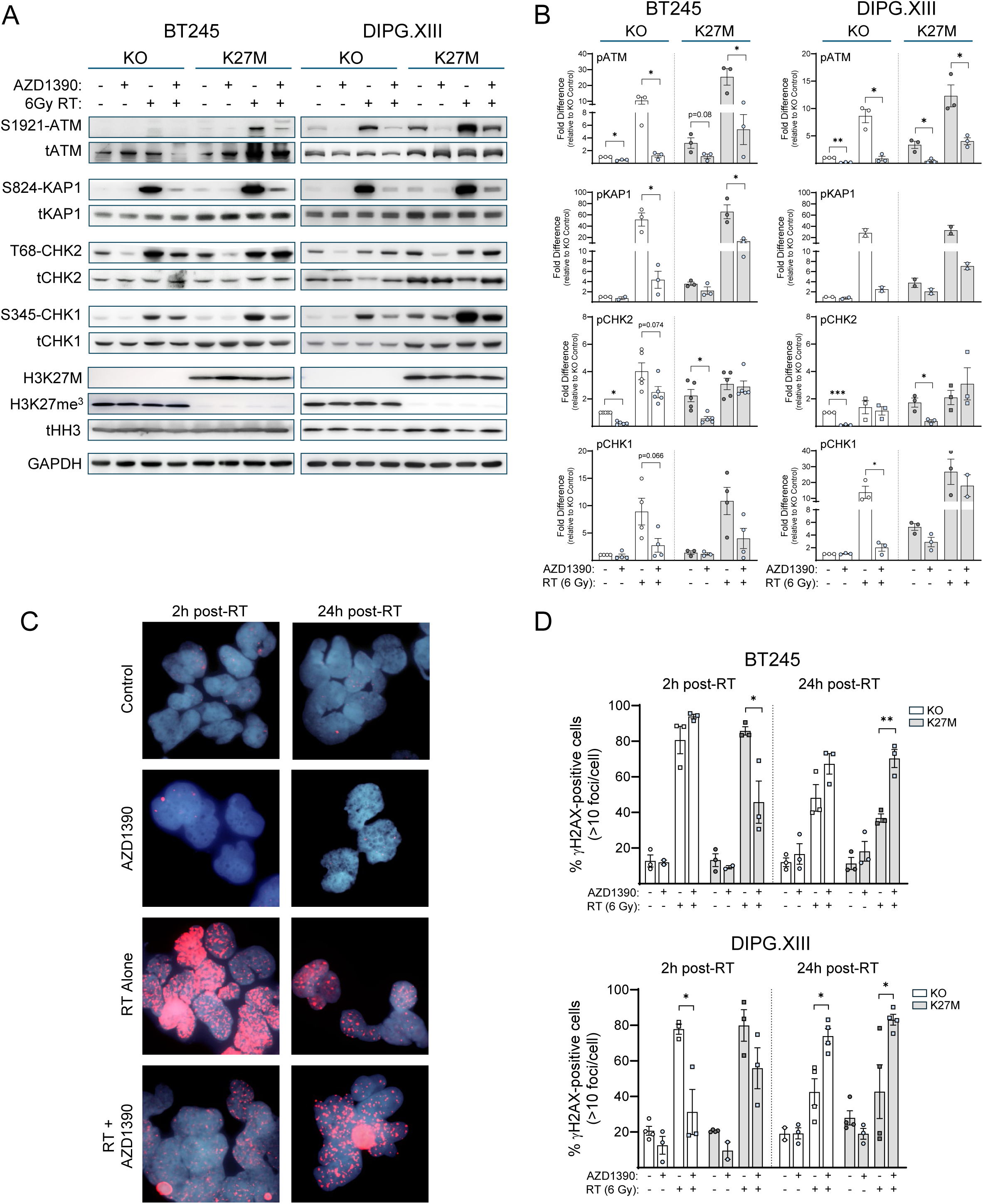
AZD1390 inhibits ATM signaling and repair of RT-induced DNA damage in DMG cells. (**A**) Immunoblots of isogenic cells treated with AZD1390 (300 nM) and RT (6 Gy) at 2 h post-RT. Images shown are representative of 4-6 independent experiments. (**B**) Quantitation of western blot images. Data are the mean ± SEM (*p<0.05). (**C**) Representative γH2AX foci in DIPG.XIII cells treated with 300 nM AZD1390. (**D**) Quantification of γH2AX foci in isogenic DMG cells 2 or 24 h post-RT. Data are the mean ± SEM (*p<0.05, **p<0.01).

To begin to understand the effects of ATM inhibition on repair of radiation-induced DNA damage in the context of DMG, we next assessed the effects of AZD1390 on radiation-induced γH2AX focus formation, as well as the persistence of γH2AX foci after 24 hours (**Fig. 4C, D**). In contrast to our finding that H3K27M-expressing cells have higher baseline levels of DNA double strand breaks (**Fig. 2C, D**) and DDR signaling (**Fig. 2A, B**), we did not see significant differences in basal γH2AX foci levels between H3K27M-expressing and H3K27M-KO cells. This difference likely reflects both redundancies in DDR signaling upstream of γH2AX, and that while the neutral comet assay specifically measures DSBs, γH2AX foci mark a variety of DNA lesions, including stalled replication forks and replication stress (30). Furthermore, differences in the ability of AZD1390 to inhibit either radiation-induced γH2AX foci at the early timepoint (2 h), or resolution of γH2AX foci at the later timepoint (24 h), did not correlate with H3K27M expression. Consistent with the ability of AZD1390 to radiosensitize both H3WT and H3K27M mutant cells, AZD1390 inhibited resolution of γH2AX foci over time in both isogenic pairs. These results support the hypothesis that AZD1390-mediated inhibition of ATM radiosensitizes DMG cells by inhibiting repair of radiation-induced DNA damage.

### Combination treatment with AZD1390 and RT is efficacious and tumor cell selective *in vivo*

We next sought to determine whether radiosensitization by AZD1390 would translate to a therapeutic benefit *in vivo*. Using a stereotactic injection method, human DIPGXIII cells harboring a spontaneous H3K27M mutation and expressing GFP and firefly luciferase (DIPG.XIII-GFP/LUC) were implanted into the cortex of Rag1-KO immune deficient mice. Animals with established tumors visualized by bioluminescent imaging (BLI; approximate flux value >1.0×10^5^ p/sec/cm^2^/sr) were randomized and treated with vehicle, AZD1390, radiation, or the combination as illustrated (**Fig. 5A**) and monitored for survival and BLI flux as primary and secondary endpoints, respectively. We found that mice treated with either AZD1390 or RT alone exhibited a modest reduction in mean BLI relative to control mice that was further inhibited by combined treatment with AZD1390 and RT (**Fig. 5B, C**). Consistent with the reduction in BLI, we observed a significant increase in survival following treatment with either AZD1390 or RT alone, resulting in a prolongation of median overall survival by 5 or 8 days, respectively, relative to vehicle control treated mice (**Fig. 5D**). Furthermore, the combination of AZD1390 with radiation significantly extended survival relative to all other treatment groups, producing a 19-day median survival benefit relative to vehicle treated mice. Treatment with AZD1390 and RT was well tolerated as reflected by minimal weight loss during therapy (**Suppl. Fig. S2**).

**Figure 5:**
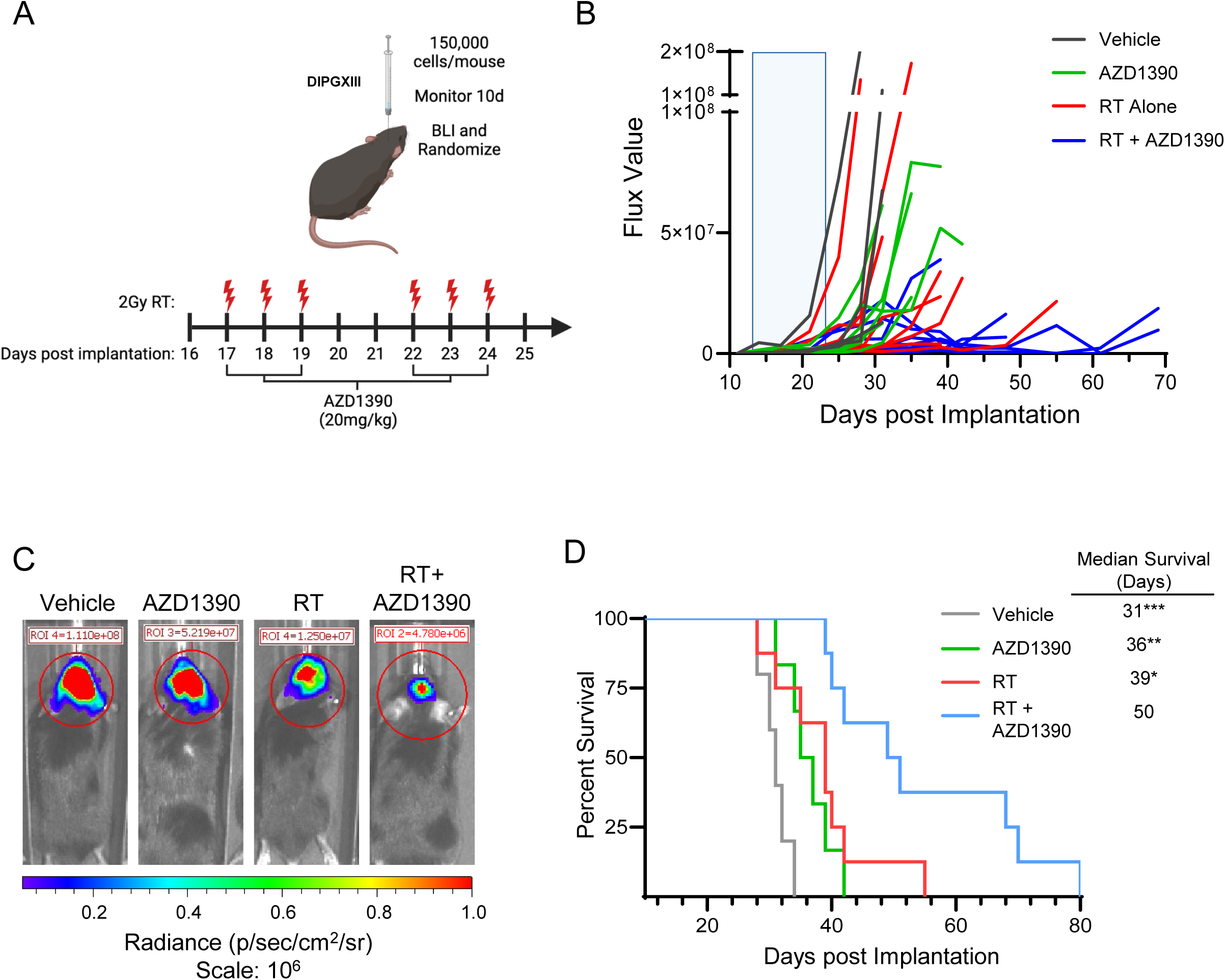
AZD1390 radiosensitizes orthotopic H3K27M DMG xenografts in vivo. (**A**) Schematic of treatment schedule, created using BioRender.com. Rag1-KO mice bearing cortical DIPG.XIII-GFP/LUC tumors were administered 20 mg/kg AZD1390 1 hour prior to each fractionated dose of radiation (red bolts) as shown. (**B**) Spider plot of bioluminescent signal flux values for individual DIPGXIII-GFP/LUC mouse tumors. Blue box indicates the treatment period. (**C**) Representative bioluminescent signal images for one mouse per treatment group, 31 or 39 days after implantation for control or treatment groups, respectively. (**D**) Kaplan-Meier survival analysis of mice bearing treated and untreated DIPGXIII-GFP/LUC orthotopic xenograft tumors (5-8 mice per treatment group).

We next conducted pharmacodynamic studies to assess the ability of AZD1390 to inhibit ATM and selectively inhibit repair of radiation-induced DNA damage, marked by persistent γH2AX staining, in DIPGXIII orthotopic tumors (31–33). Tumor cells, identified by GFP expression, and adjacent normal brain were harvested from mice on the second day of treatment (day 18 as illustrated in **Fig. 5A**) and AZD1390 activity was assessed by either immunoblotting for pATM (S1981) (**Fig. 6A, B**) or pATM IHC staining (**Fig. 6C, D**). As anticipated, RT-mediated pATM was significantly inhibited by AZD1390, confirming the high penetrance of AZD1390 into orthotopic brain tumors. We next evaluated γH2AX levels in both tumor and adjacent normal brain to determine whether inhibition of ATM caused persistent radiation-induced DNA damage *selectively* in DIPGXIII tumors (**Fig. 6C, E**). γH2AX staining trended to be higher in tumor versus normal tissue, and was attenuated by AZD1390 in tumor but not in normal brain (though this effect did not reach statistical significance). Taken together, these data demonstrate effective AZD1390-mediated inhibition of ATM in an orthotopic model of DMG, and suggest the effects of AZD1390 on repair of RT-induced DNA damage are tumor-cell selective, relative to adjacent normal brain tissue.

**Figure 6:**
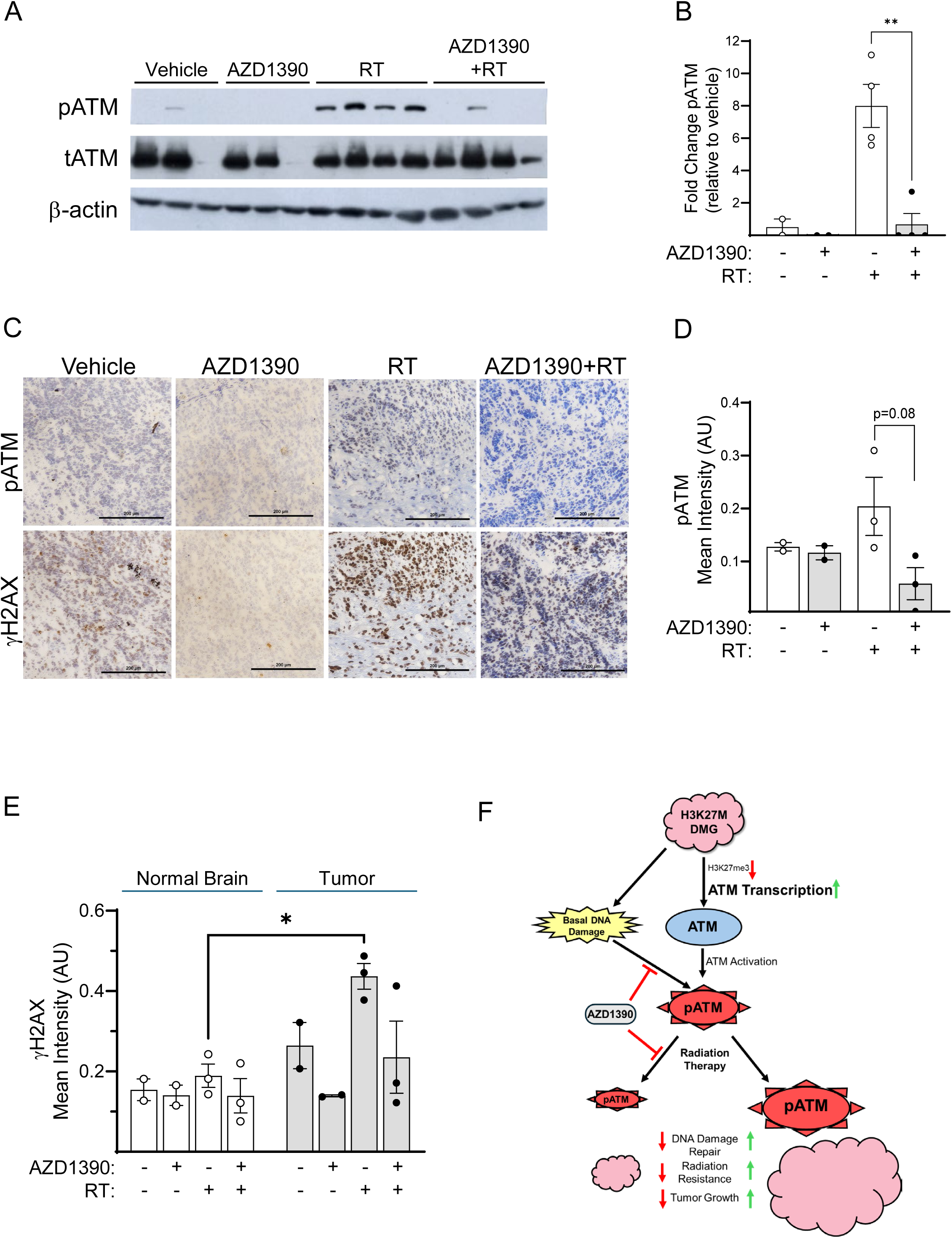
AZD1390 inhibits radiation-induced ATM activity and induces persistent DNA damage in H3K27M DMG xenografts. (**A**) Immunoblots of orthotopic H3K27M mutant DIPGXIII xenografts harvested on day 2 of treatment (18 days post-implantation) as illustrated in Fig. 5A. Samples are from 3 – 4 independent tumors for each condition. (**B**) Quantitation of pATM (S1981) immunoblots. Data are the mean ± SEM with statistical significance indicated **p<0.01. (**C**) Representative images of immunohistochemistry for pATM (S1981) and γH2AX at day 18 from orthotopic H3K27M mutant DIPGXIII xenografts. (**D**) Quantitation of pATM (S1981) immunohistochemistry. Data are the mean ± SEM DAB intensity of 2-3 independent tumors. Five regions of interest from 3 fields were scored for each tumor. (**E**) Quantitation of γH2AX immunohistochemistry in both orthotopic tumors and adjacent normal brain. Data are the mean ± SEM DAB intensity of 2-3 independent samples with 5 regions of interest from 3 fields scored for each tissue and statistical significance indicated *p<0.05. (**F**) Model depicting the effects of AZD1390 on constitutive and RT-mediated ATM signaling in the context of H3K27M DMG.

## DISCUSSION

In this study, we have demonstrated a direct mechanistic link between the H3K27M mutation that drives DMG tumorigenesis and the apical DNA damage response kinase ATM (**Fig. 6F**). We found that cells expressing the histone H3K27M protein have reduced levels of the transcription repressor mark H3K27me^3^ in the *ATM* promoter, an epigenetic change that was associated with increased ATM expression and signaling activity. While H3K27M-expressing cells have higher levels of basal DNA damage, we did not see striking differences in viability or radiosensitivity between the isogenic lines. These results suggest that ATM hyperactivity in H3K27M-expressing cells may promote survival in the presence of high levels of genomic instability. Furthermore, we show that the H3K27M tumor-specific ATM hyperactivity can be therapeutically leveraged to selectivity treat tumor cells while avoiding toxicity in surrounding normal tissue.

Our findings contribute to the expanding field of research showing that DMG tumors present therapeutic vulnerabilities in their dependence on DNA damage repair pathways. Several groups have explored targeting DDR proteins to enhance radiation efficacy in pHGG, with efforts focused on inhibiting WEE1, CHK1/2 or altered TP53 states, in addition to ATM (11,34,35). An understanding, however, of whether the H3K27M mutation directly and/or specifically influences either radiosensitivity or susceptibility to ATM inhibitor-mediated radiosensitization was previously unclear. In the isogenic models used in our study, we found that while H3K27M knock-out (H3K27M-KO) attenuated constitutive ATM activity and reduced basal levels of DNA DSBs, it did not consistently increase radiosensitivity. Furthermore, while there was a trend towards increased AZD1390-mediated radiosensitization in the genetically engineered isogenic murine model of H3K27M-expressing DMG relative to H3K27 wild-type (Fig. 3A; p=0.064), H3K27M-KO in patient-derived DMG cell lines did not significantly decrease AZD1390-mediated radiosensitization. Several factors that may contribute to this result, including additional mutations in the patient derived xenografts that confer sensitivity to ATM inhibition but are not affected by loss of H3K27M or the relatively high (though still clinically achievable) concentration of AZD1390 used in our studies. However, given the critical role of ATM in the DNA damage response and the body of literature documenting the radiosensitizing activity of ATM inhibition in a variety of tumor models, the ability of AZD1390 to radiosensitize DMG cells lacking the H3K27M mutation is not surprising. Furthermore, as not all pHGG express the H3K27M mutation, this susceptibility could be therapeutically beneficial.

While the results of these studies are promising, unanswered questions remain. For example, given that H3K27M causes global de-repression of gene expression, it is unclear why ATM specifically is upregulated in H3K27M tumors relative to other DDR kinases. Furthermore, it is not known whether ATM is upregulated, or whether combined treatment with ATM inhibitor and RT is efficacious, in either pHGG with different histone H3 mutations (eg, H3G34R), or tumors such as posterior fossa ependymoma that have *H3K27M-like* alterations (eg, EZHIP overexpression) similarly associated with global reprogramming of the epigenome (36). In addition, we do not know whether the H3K27M mutation directly affects recruitment of ATM or other DDR proteins to radiation-induced DSBs. One of the global epigenetic changes associated with the H3K27M mutation is increased H3K36 dimethylation (H3K36me^2^), an epigenetic mark that serves as a docking sight for the DNA repair complex MRN (MRE11, RAD50, and NBS1) and subsequent recruitment and activation of ATM (37,38). ATM-mediated phosphorylation of lysine-specific demethylase 2A (KDM2A) attenuates the ability of KDM2A to bind chromatin, preserving H3K36me^2^ at sites of damage and promoting DNA repair (37,39–41). It is possible that H3K27M mutant cells harness these actions to create a DNA repair positive feedback loop that mitigates the effects of radiation on tumor cell viability, but this hypothesis requires further investigation.

One of the primary concerns for the clinical translation of ATM inhibitors as radiosensitizing agents for patients with pHGG is the development of radiation-induced brain injury. Thus far, however, preclinical studies have found little evidence of durable CNS toxicity in mice treated with AZD1390 and directed radiation therapy (33,34). Furthermore, recently presented results from an ongoing Phase I clinical trial (NCT03423628) suggest AZD1390 in combination with intensity-modulated radiation therapy (IMRT) has an acceptable safety profile and therapeutic index in adult patients with brain malignancies (35). Our results (Fig. 6D,E) suggest that AZD1390 minimally affects RT-induced ATM signaling in non-cancerous brain, further supporting the existence of a favorable therapeutic index for this drug.

In conclusion, our findings have important implications for the development of more effective DMG treatment strategies. Most patients with DMG receive radiation as monotherapy. Our promising preclinical data, combined with additional published data showing that AZD1390 improves the efficacy of radiation, suggest that combination studies with ATM inhibition and radiation should be explored (9,31–33). The tumor selectivity seen in our study and the and clinical safety of combined radiation and AZD1390 in adult brain tumor patients suggests that this approach will be well tolerated (33). Collectively, these findings support the further clinical development of AZD1390 with radiotherapy as a therapeutic strategy for H3K27M mutant pediatric gliomas like DMG.

## Acknowledgements

This work was supported by R01CA240515 (M.A.M.); U01CA216449 (T.S.L.); P50CA269022 (M.A.M. and T.S.L.), R50CA251960 (L.A.P.), the Chad Carr Pediatric Brain Tumor Center (M.A.M., Q.Z. and E.P.), the SPORE Career Enhancement Program (Q.Z.), Alex’s Lemonade Stand (M.A.M., D.R.W.), National Institutes of Health Grants R01-NS119231, R01-NS124607, DOD Grant CA201129P1 (C.K.), and the Rogel Comprehensive Cancer Center (P30CA046592). D.R.W. is also supported by the NCI (K08CA234416; R37CA258346), the NINDS (R01NS129123), the V Foundation, the Damon Runyon Cancer Foundation, the Sontag Foundation, the Ivy Glioblastoma Foundation, the Forbes Institute for Cancer Discovery, and the Chad Tough Defeat DIPG foundation.

## Acknowledgements

The authors would like to thank Dr. Nada Jabado for providing the BT245 and DIPG.XIII parental and H3K27M-KO isogenic cells lines.

## SUPPLEMENTARY FIGURE LEGENDS

**Table S1:**
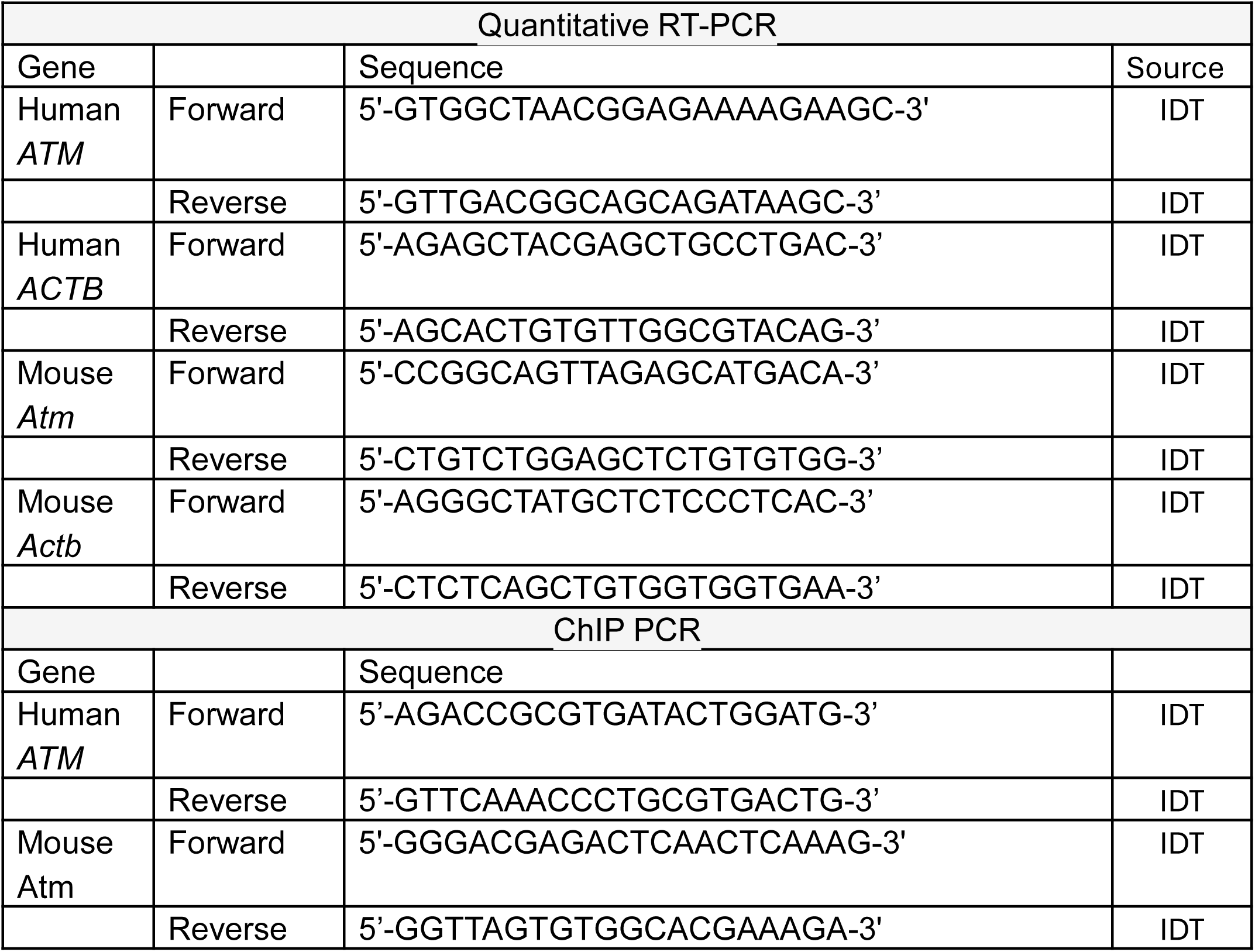
Primers used for Quantitative RT-PCR and ChIP PCR analysis.

**Table S2:** Antibodies used for ChIP PCR, immunoblotting, immunofluorescence staining and immunohistochemistry.

| ChIP PCR |  |  |  |  |
| --- | --- | --- | --- | --- |
| Target | Clone | Species | Catalogue Number | Source |
| Normal Rabbit IgG |  | rabbit | 2729 | Cell Signaling |
| Total Histone H3 | D2B12 | rabbit | 4620 | Cell Signaling |
| H3K27me<sup>3</sup> |  | rabbit | 07-449 | Millipore Sigma |
| Western Blot Analyses |  |  |  |  |
| Target | Clone | Species | Catalogue Number | Source |
| Total ATM | D2E2 | rabbit | 2873 | Cell Signaling |
| pATM (S1981) | D25E5 | rabbit | 13050 | Cell Signaling |
| Total KAP1 | C42G12 | rabbit | 4124 | Cell Signaling |
| pKAP1 (S824) |  | rabbit | 4127 | Cell Signaling |
| *Total CHK2 | N-17 | goat | sc-8812 | Santa Cruz Biotechnology |
| Total CHK2 | Clone 7 | mouse | 05-649 | Millipore Sigma |
| pCHK2 (T68) | C13C1 | rabbit | 2197 | Cell Signaling |
| Total CHK1 | 2G1D5 | mouse | 2360 | Cell Signaling |
| pCHK1 (S345) | 133D3 | rabbit | 2348 | Cell Signaling |
| Total Histone H3 |  | rabbit | 9715 | Cell Signaling |
| Histone H3<br>(K27M-specific) | D3B5T | rabbit | 74829 | Cell Signaling |
| Trimethyl Histone H3<br>(K27) | C36B11 | rabbit | 9733 | Cell Signaling |
| GAPDH | 14C10 | rabbit | 2118 | Cell Signaling |
| β-actin | C4 | mouse | sc-47778 | Santa Cruz Biotechnology |
| Immunofluorescence Staining |  |  |  |  |
| Target | Clone | Species | Catalogue Number | Source |
| γH2AX | JBW301 | mouse | 05-636 | Millipore Sigma |
| Immunohistochemistry |  |  |  |  |
| pATM (S1981) | D25E5 | rabbit | 13050 | Cell Signaling |
| γH2AX | JBW301 | rabbit | 2577 | Cell Signaling |

\*This antibody is no longer commercially available. Similar results were obtained with the alternate total CHK2 antibody listed above.

**Table S3:**
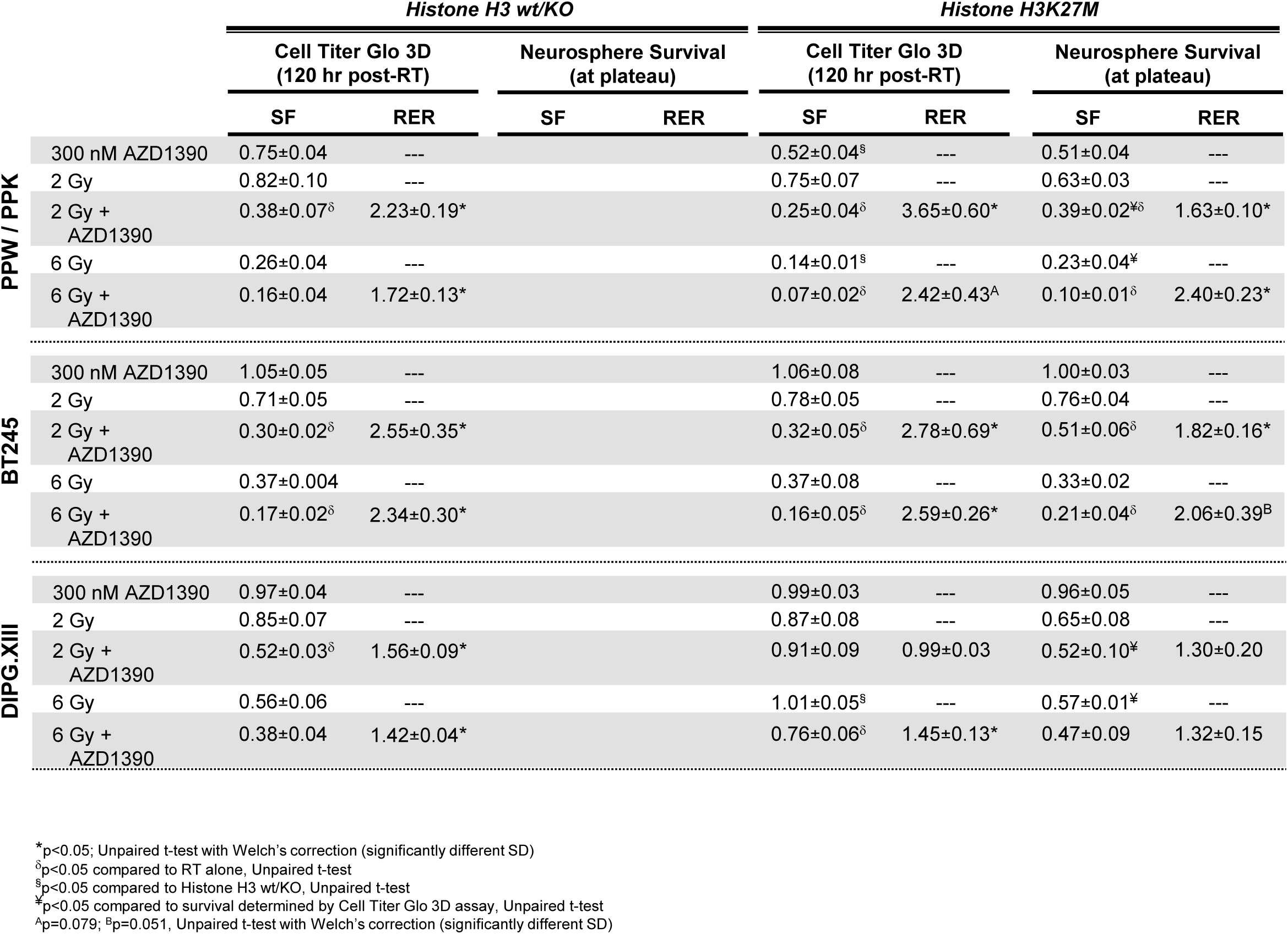
Cytotoxicity and radiosensitization by ATM inhibition in isogenic H3K27M DMG cell lines.

**Figure S1:**
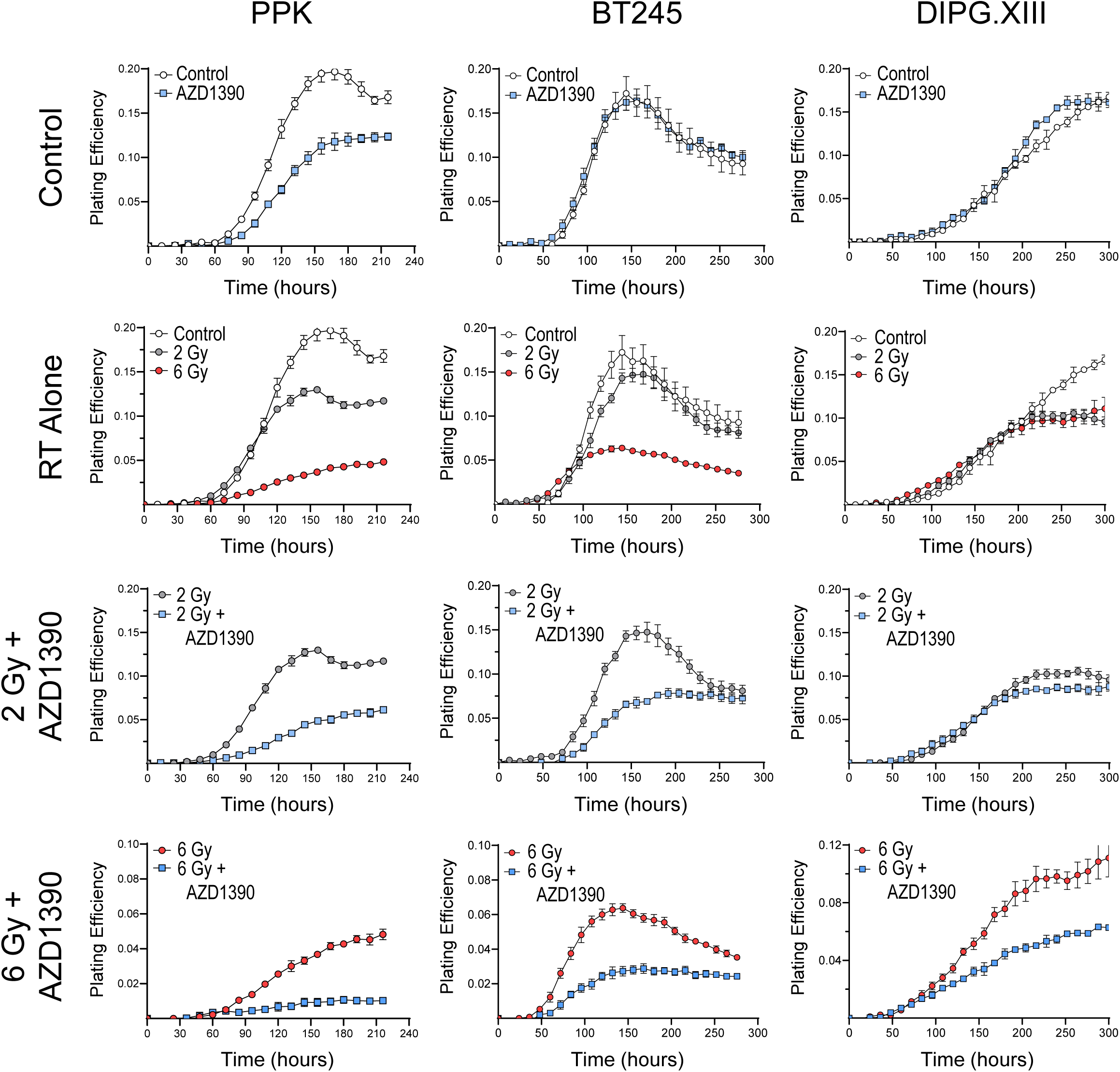
The effects of AZD1390 and radiation on neurosphere growth over time as assessed with live-cell imaging. Data are the mean ± SEM of technical quaduplicates from a single representative experiment for each cell line. For experimental details, see *Methods*.

**Figure S2:**
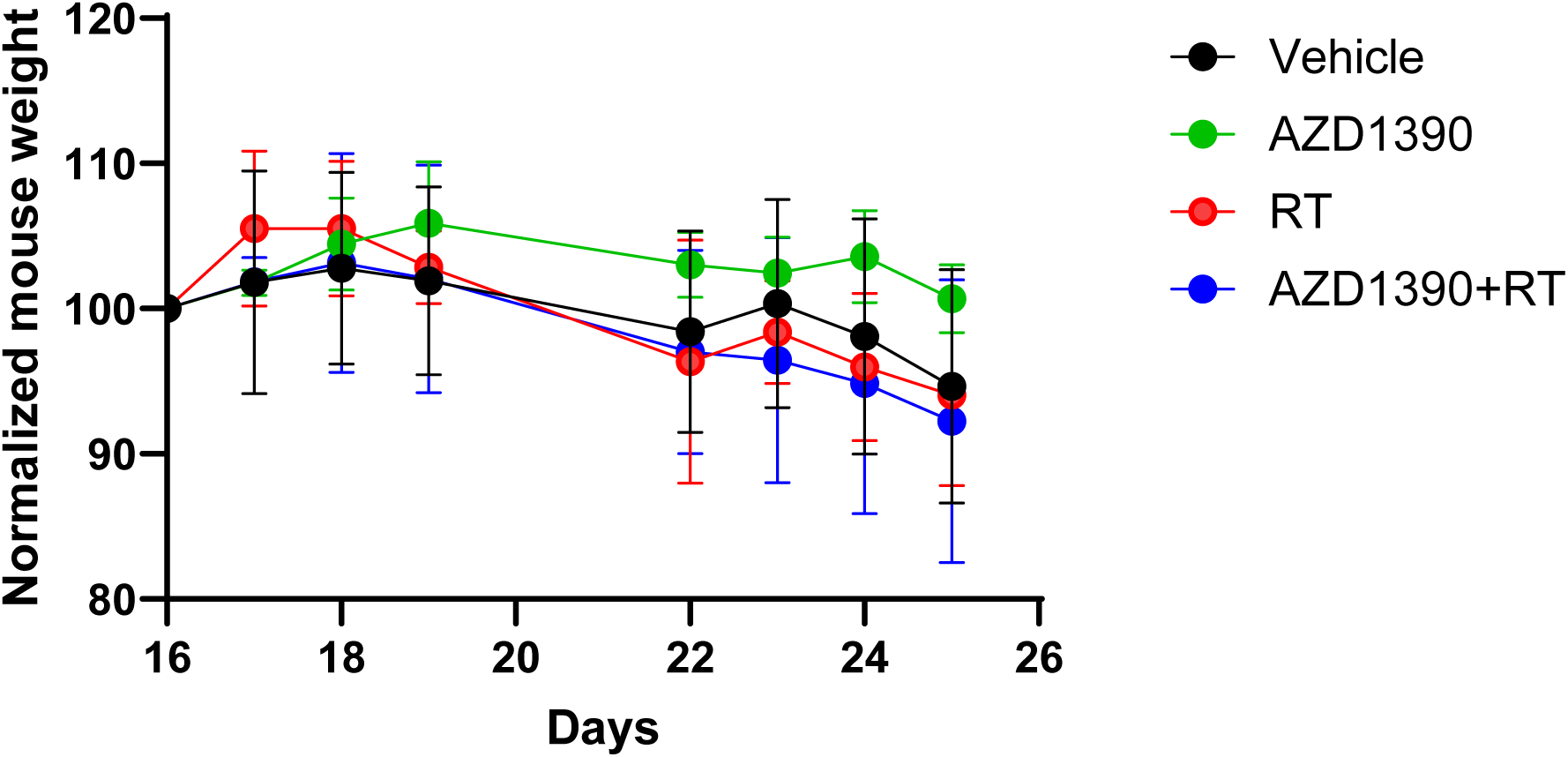
The effects of AZD1390 and radiation on mouse weight during therapy. Mouse weights were normalized to body weight on Day 0. Data are the mean ± SEM (n=5-8).

## Notes

**Conflict-of-interest disclosure:** M.A.M. has received research funding and honoraria from AstraZeneca.

### Competing Interest Statement

M.A.M. has received research funding and honoraria from AstraZeneca.

